# Gains and Losses affect Learning Differentially at Low and High Attentional Load

**DOI:** 10.1101/2020.09.01.278168

**Authors:** Kianoush Banaie Boroujeni, Marcus Watson, Thilo Womelsdorf

## Abstract

Prospective gains and losses modulate cognitive processing, but it is unresolved whether gains and losses can facilitate flexible learning in changing environments. The prospect of gains might enhance flexible learning through prioritized processing of reward-predicting stimuli but is unclear how far this learning benefit extends when task demands increase. Similarly, experiencing losses might facilitate learning when they trigger attentional re-orienting away from loss-inducing stimuli, but losses may also impair learning by reducing the precise encoding of loss-inducing stimuli. To clarify these divergent views, we tested how varying magnitudes of gains and losses affect the flexible learning of object values in environments that varied attentional load by increasing the number of interfering object features during learning. With this task design we found that larger prospective gains improved learning efficacy and learning speed, but only when attentional load was low. In contrast, expecting losses generally impaired learning efficacy and this impairment was larger at higher attentional load. These findings functionally dissociate the contributions of prospective gains and losses on flexible learning, suggesting they operate via separate control mechanisms. One process is triggered by experiencing loss and seems to disrupt the encoding of specific loss-inducing features which leads to less efficient exploration during learning. The second process is triggered by experiencing gains which enhances learning through a more efficient prioritizing of reward-predicting stimulus features as long as the interference of distracting information is limited. These results demonstrate strengths and limitations of motivational regulation of learning efficacy in multidimensional environments having variable attentional loads.

**Significance statement:** Increasing the prospective gains is assumed to enhance flexible learning, but there is no consensus on whether imposing losses enhances or impairs flexible learning. We show that anticipating loss of already attained assets generally reduced learning changes in the relevance of visual objects and that this learning impediment is more pronounced when learning demands higher attentional control of interference from distracting object features. Moreover, we show that increasing the prospective gains indeed facilitates learning, but only when the learning problem has intermediate or low attentional demands. These findings document that the beneficial effects of gains hit a limit when task demands increase, and that prospective losses reduce cognitive flexibility already at low task demands which is exacerbated when task demands increase. These findings provide novel insight into the strengths and limitations of gains and of losses to support flexible learning in multidimensional environments imposing variable attentional loads.

## Introduction

Anticipating gains or losses have been shown to enhance the motivational saliency of information (Berridge & Robinson, 2016; Failing & Theeuwes, 2018; Yechiam & Hochman, 2013b). Enhanced motivational saliency can be beneficial when learning about the behavioral relevance of visual objects. Learning which objects lead to higher gains should enhance the likelihood choosing those objects in the future to maximize rewards, while learning which objects lead to loss should enhance avoiding those objects in the future to minimize loss (Collins & Frank, 2014; Maia, 2010). While these scenarios are plausible from a rational point of view, empirical evidence suggests more complex consequences of gains and losses for adaptive learning.

Prospective gains are generally believed to facilitate learning and attention to reward predictive stimuli. But most available evidence is based on tasks using simple stimuli, leaving open whether the benefit of gains generalizes to settings with more complex multidimensional objects that have high demands on cognitive control. With regard to losses, there is conflicting evidence with some studies showing benefits and others showing deterioration of performance when subjects experience or anticipate losses. It is not clear whether these conflicting effects of loss are due to a u-shaped dependence of loss effects on performance with only intermediate levels having positive effects (Yechiam, Ashby, & Hochman, 2019; Yechiam & Hochman, 2013a), or whether experiencing loss reduces the encoding of loss-inducing events in favor of a more generalized re-orientation away from loss-inducing situations (Barbaro, Peelen, & Hickey, 2017; Laufer, Israeli, & Paz, 2016; McTeague, Gruss, & Keil, 2015). To shed light on the impact of gains and losses for flexible learning does therefore require an experiment that varies not only the degree of gains and losses during learning to possibly discern a u-shaped effect on learning, but that also varies the attentional demands of the learning task to potentially see the limitations of prospective gains and losses in supporting flexible learning. This study details such an experiment to identify how gains and losses improve or impair flexible learning at low and high attentional demands.

It is widely believed that prospective gains improve attention to reward predicting stimuli, which predicts that learning about stimuli should be facilitated by increasing their prospective gains. According to this view anticipating gains acts as an independent ‘top-down’ mechanism for attention, which has been variably described as *value-based attentional guidance* (Anderson, 2019; Bourgeois, Chelazzi, & Vuilleumier, 2016; Theeuwes, 2019; Wolfe & Horowitz, 2017), *attention for liking* (Gottlieb, 2012; Hogarth, Dickinson, & Duka, 2010), or *attention for reward* (San Martin, Appelbaum, Huettel, & Woldorff, 2016). These attention frameworks suggest that stimuli are processed quicker and with higher accuracy when they become associated with positive outcomes (Barbaro et al., 2017; Hickey, Kaiser, & Peelen, 2015; Schacht, Adler, Chen, Guo, & Sommer, 2012). The effect of anticipating gains can be adaptive when the gain associated stimulus is a target for goal-directed behavior, but it can also deteriorate performance when the gain-associated stimulus is distracting or task irrelevant in which case it attracts attentional resources away from more relevant stimuli (Chelazzi, Marini, Pascucci, & Turatto, 2019; Noonan, Crittenden, Jensen, & Stokes, 2018).

Compared to gains, the behavioral consequences of experiencing or anticipating loss are often described in affective and motivational terms rather than in terms of attentional facilitation or impediment (Dunsmoor & Paz, 2015). An exception to this is the so-called ‘*loss attention’* framework that describes how losses trigger shifts in attention to alternative options and thereby improve learning outcomes (Yechiam & Hochman, 2013a, 2014). The *loss attention* framework is based on the finding that experiencing loss causes a vigilance effect that triggers enhanced exploration of alternative options (Lejarraga & Hertwig, 2017; Yechiam & Hochman, 2013a, 2013b, 2014). The switching to alternative options following aversive loss events can be observed even when the expected values of the available alternatives are controlled for and is not explained away by an affective loss aversion response as subject with higher or lower loss aversion both show loss-induced exploration (Lejarraga & Hertwig, 2017). According to these insights, experiencing loss should improve adaptive behavior by facilitating avoidance of bad options. Consistent with this view humans and monkeys have been shown to avoid looking at visual objects paired with unpleasant consequences (such as a bitter taste or a monetary loss) (Ghazizadeh, Griggs, & Hikosaka, 2016b; Raymond & O’Brien, 2009; Schomaker, Walper, Wittmann, & Einhauser, 2017).

What is unclear, however, is whether loss-triggered shifts of attention away from a stimulus does allow for the precise encoding of the loss-inducing stimulus, or rather reflects an unspecific re-orienting away from a loss evoking situation. The empirical evidence about this question is contradictory with some studies reporting better encoding and memory for loss-evoking stimuli, while other studies reporting poorer memory and insights about the precise stimuli that triggered the aversive outcomes. Evidence of a stronger memory representation of aversive outcomes comes from studies reporting increased detection speed of stimuli linked to threat-related aversive outcomes such as electric shocks (Ahs, Miller, Gordon, & Lundstrom, 2013; Li, Howard, Parrish, & Gottfried, 2008), painful sounds (McTeague et al., 2015; Rhodes, Ruiz, Rios, Nguyen, & Miskovic, 2018), threat-evoking air puffs (Ghazizadeh, Griggs, & Hikosaka, 2016a), or fear-evoking images (Ohman, Flykt, & Esteves, 2001). In these studies, subjects were faster or more accurate in responding to stimuli that were associated with aversive outcomes. This improved responding to these threat-related aversive stimuli indicates that those stimuli are stronger represented than neutral stimuli and hence can guide adaptive behavior away from them. Notably, such a benefit is not restricted to threat-related aversive stimuli but can also be seen in faster response times to stimuli associated with the loss of money, which is a secondary ‘learned’ reward (Bucker & Theeuwes, 2016; Small et al., 2005; Suarez-Suarez, Holguin, Cadaueira, Nobre, & Doallo, 2019). For example, in an object-in-scene learning task attentional orienting to the incorrect location was faster when subjects lost money for the object at that location in prior encounters compared to a neutral or positive outcome (Doallo, Patai, & Nobre, 2013; Suarez-Suarez et al., 2019). Similarly, when subjects are required to discriminate a peripherally presented target object they detect the stimulus faster following a short (20ms) spatial pre-cue when the cued stimulus is linked to a monetary loss (Bucker & Theeuwes, 2016). This faster detection was similar for monetary gains indicating that gains and losses can have near-symmetric beneficial effects on attentional capture. A similar benefit for loss-as well as gain-associated stimuli has also been reported when stimuli are presented briefly and subjects have to judge whether the stimulus had been presented before (O’Brien & Raymond, 2012). The discussed evidence suggests that loss-inducing stimuli have a processing advantage for rapid attentional orienting and fast perceptual decisions even when the associated loss is a secondary reward like money. However, whether this processing advantage for loss associated stimuli can be used to improve flexible learning and adaptive behavior is unresolved.

An alternate set of studies contradicts the assumption that experiencing loss may improve flexible behavior by investigating not the rapid orienting away from aversive stimuli but the fine perceptual discrimination of stimuli associated with negative outcomes (Laufer et al., 2016; Laufer & Paz, 2012; Resnik, Laufer, Schechtman, Sobel, & Paz, 2011; Schechtman, Laufer, & Paz, 2010; Shalev, Paz, & Avidan, 2018). In these studies, anticipating the loss of money did not enhance but systematically reduced the processing of loss-associated stimuli, causing impaired perceptual discriminability and reduced accuracy, even when this implied losing money during the experiment (Barbaro et al., 2017; Laufer et al., 2016; Laufer & Paz, 2012; Shalev et al., 2018). For example, losing money for finding objects in natural scenes reduces the success rate of human subjects to detect those objects compared to searching for objects that promises gains (Barbaro et al., 2017). One important observation in these studies is that the detrimental effect of losses is not simply explained away by an overweighting of losses over gains, which would be suggestive of an affective loss aversion mechanism (Laufer & Paz, 2012). Rather, this literature suggests that stimuli linked to monetary loss outcomes are weaker attentional targets compared with neutral stimuli or gain-associated stimuli. This weaker representation can be found with multiple types of aversive outcomes including when stimuli are associated with monetary loss (Laufer et al., 2016; Laufer & Paz, 2012), unpleasant odors (Resnik et al., 2011), or electric shock (Shalev et al., 2018). One possible mechanism underlying this weakening of stimulus representations following aversive experience is that aversive outcomes are less precisely linked to the stimulus causing the outcome. According to this account, the credit assignment of an aversive outcome might generalize to other stimuli that share attributes with the actual stimulus whose choice caused the negative outcome. Such a generalized assignment of loss can have positive as well as negative consequences for behavioral performance (Dunsmoor & Paz, 2015; Laufer et al., 2016). It can lead to faster detection of loss-threatening stimuli in similar situations to the original aversive situation without requiring recognizing the precise object instance that was causing the initial loss. This may lead to enhanced learning in situations in which the precise stimulus features are not critical. However, the wider generalization of loss outcomes will be detrimental in situations that require the precise encoding of the object instance that gave rise to loss.

In summary, the surveyed empirical evidence suggests a complex picture about how losses may influence adaptive behaviors and flexible learning. On the one hand, experiencing or anticipating loss may enhance learning, when it triggers attention shifts away from the loss-inducing stimulus and when it enhances fast recognition of loss-inducing stimuli to more efficiently avoid them. But on the other hand, evidence suggests that associating loss with a stimulus can impair learning when the task demands require precise insights about the loss-inducing stimulus features, because these features may be less well encoded after experiencing losses than gains, which will reduce their influence on behavior or attentional guidance in future encounters of the stimulus.

To understand which of these scenarios holds true, we designed a learning task that varied two main factors. First, we varied the magnitude of gains and losses to understand whether the learning effects depend on the actual gain/loss magnitudes that are only rarely manipulated in the discussed studies (above). To achieve this, we used a token system in which subjects receive tokens for correct and loose tokens for incorrect choices. Secondly, we varied the attentional demands of the learning problems by requiring subjects to explore object features in either only one dimension (e.g. different colors) to find a rewarded feature, or to explore object features in two or three dimensions (e.g. different colors, body shapes and body pattern) when searching for the rewarded feature. The variation of the object feature dimensionality allows testing whether losses and gains differentially facilitate or impair learning at varying load on attentional processing.

With this task design we found across four rhesus monkeys first, that expecting larger gains enhanced the efficacy of learning targets particularly at low attentional load, but not at the highest attentional load. Secondly, we found that experiencing loss generally decreased flexible learning, that larger losses exacerbate this effect, and that the loss induced learning impairment is worse at high attentional load.

Our study uses nonhuman primates as subjects to establish a robust animal model for understanding the influence of gains and losses on learning in a task with translational value for humans as well as other species. Establishing this animal model will facilitate future studies about the underlying neurobiological mechanisms. Leveraging this animal model is possible because nonhuman primates readily understand a token reward/punishment system similar to humans and can track sequential gains and losses of assets before cashing them out for primary (e.g. juice) rewards (Rich & Wallis, 2017; Seo & Lee, 2009; Shidara & Richmond, 2002; Taswell, Costa, Murray, & Averbeck, 2018).

## Materials and Methods

### Experimental Procedures

All animal related experimental procedures were in accordance with the National Institutes of Health Guide for the Care and Use of Laboratory Animals, the Society for Neuroscience Guidelines and Policies, and approved by the Vanderbilt University Institutional Animal Care and Use Committee.

Four pair-housed male macaque monkeys (8.5-14.4 kg, 6-9 years of age) were allowed separately to enter an apartment cage with a custom build, cage-mounted touchscreen Kiosk-Cognitive-Engagement Station to engage freely with the experimental task for 90-120 min per day. The Kiosk-Cognitive-Engagement Station substituted the front panel of an apartment cage with a 30 cm recessed, 21’ touchscreen and a sipper tube protruding towards the monkey at a distance of ∼33 cm and a height that allows all animals sitting comfortably in front of the sipper tube with the touchscreen in reaching distance. Details about the Kiosk Station and the training regime are provided in (Womelsdorf et al., in preparation). In brief, all animals underwent the same training regimes involving learning to touch, hold and release touch in a controlled way at all touchscreen locations. Then animals learned visual detection and discrimination of target stimuli among increasingly complex non-rewarded distracting objects, with correct choices rewarded with fluid. Objects were 3-dimensionally rendered Quaddles that have a parametrically controllable feature space, varying along four dimensions, including the color, body shape, arm types and surface pattern (Watson, Voloh, Naghizadeh, & Womelsdorf, 2019). Throughout training, one combination of features was never rewarded and hence termed ‘neutral’, which was the spherical, uniform, grey Quaddle with straight, blunt arms (**Fig. 1C**). Relative to the features of the neutral object we could then increase the feature space for different experimental conditions by varying from trial-to-trial features from only one, two, or three dimensions relative to the neutral object. After monkeys completed the training for immediate fluid reward upon a correct choice, a token history bar was introduced and animals performed the feature learning task for receiving tokens until they filled the token history bar with five tokens to cash out for fluid reward (**Fig. 1A**). All animals effortlessly transitioned from immediate fluid reward delivery for correct choices to the token-based reward schedules.

**Figure 1.**
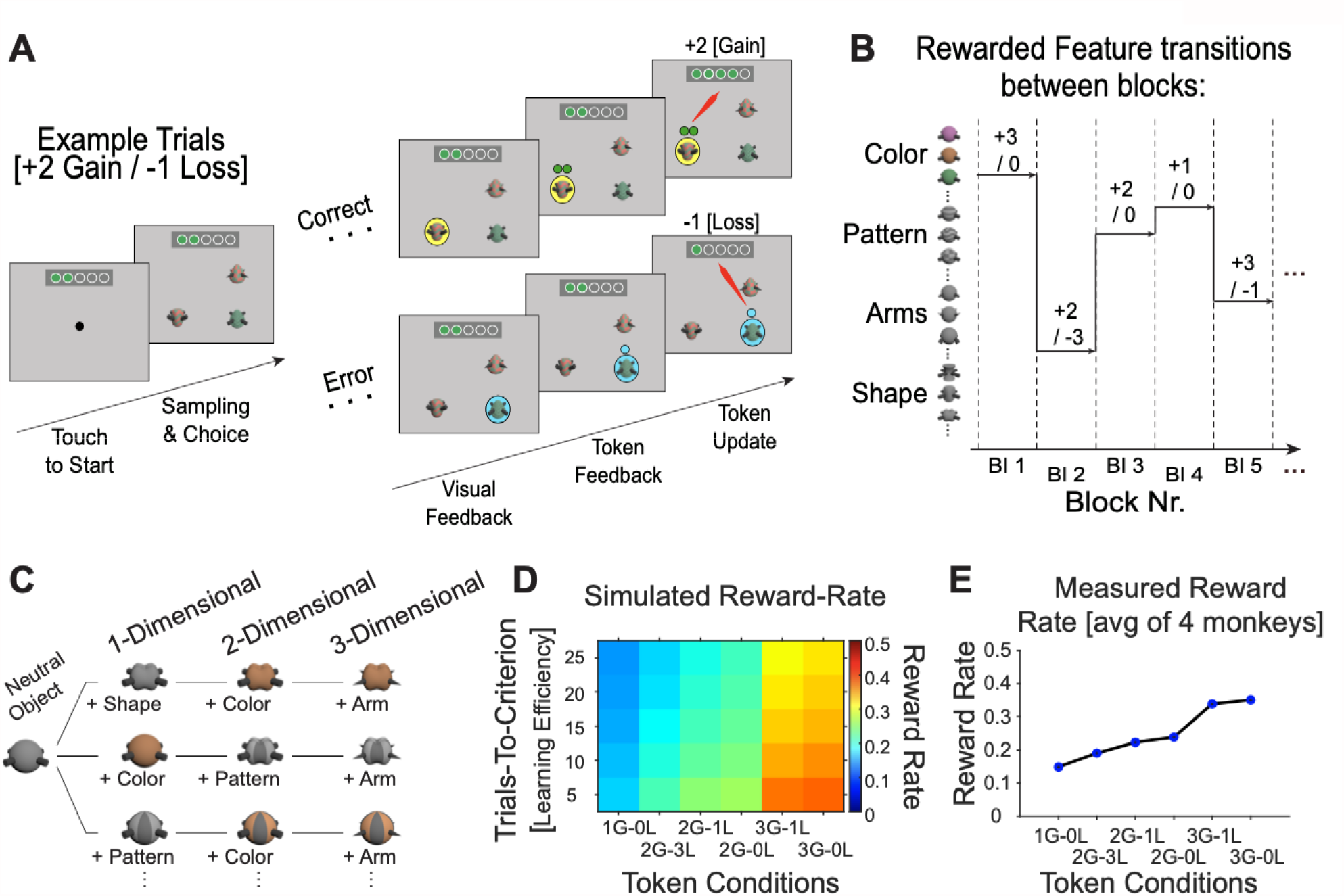
Task paradigm varying attentional load and token gains and losses. **A**: The trial sequence starts with presenting 3 objects. The monkey choses one by touching it. Following a correct choice a yellow halo provides visual feedback, then green tokens are presented above the stimulus for 0.3s before they are animated to move upwards towards the token bar where the gained tokens are added. Following an error trial visual feedback is cyan, then empty token(s) indicate the amount of tokens that will be lost (here: -1 token). The token update moves the empty token to the token bar where green tokens are removed from the token bar. When ≥5 tokens have been collected the token bar blinks red/blue, three drops of fluid are delivered, and the token bar reset. **B:** Over blocks of 35-60 trials one object feature (here: in rows) is associated with either 1,2, or 3 token gains, while objects with other features are linked to either, 0, -1, or -3 token loss. **C:** Attentional load is varied by increasing the number of features that define objects. The one-dimensional (1D) load condition presents objects that vary in only one feature dimension relative to a neutral object, the 2D load varies features of two feature dimensions, and the 3D load condition varies three feature dimensions. For example, a 3D object has different shapes, colors, and arm types across trials (but the same neutral pattern). **D:** Simulation of the expected reward rate animals receive with different combinations of token-gains and -losses (*x-axis*) given different learning speed of the task (*y-axis*). **E:** Actual reward rates (y-axis) for different token conditions (*x-axis*) based on their learning speed across 4 monkeys.

The visual display, stimulus timing, reward delivery and registering of behavioral responses was controlled by the Unified Suite for Experiments (USE), which integrates an IO-controller board with a unity3D video-engine based control for displaying visual stimuli, controlling behavioral responses, and triggering reward delivery (Watson, Voloh, Thomas, Hasan, & Womelsdorf, 2019).

### Task Paradigm

The task required monkeys to learn a target feature in blocks of 35-60 trials by choosing one among three objects, each composed of multiple features (**Fig. 1A-C**). At the beginning of each trial, a blue square with a side length of 3.5 cm (3° radius wide) appeared on the screen. To start a new trial, monkeys were required to touch and hold the square for 500 ms. Within 500 ms after the blue square touch was accepted, three stimuli appeared on the screen randomly on 3 out of 4 possible locations with an equal distance from the screen center (10.5 cm, 17° eccentricity). Each stimulus had a diameter of 3 cm (∼2.5° radius wide) and was horizontally and vertically distanced from other stimuli by 15 cm (24°). To select a stimulus, monkeys were required to touch the object for a duration longer than 100 ms. If a touch was not registered within 5 seconds after the appearance of the stimuli, the trial was aborted.

Each experimental session consisted of 36 learning blocks. Each learning blocks included 35-60 trials. Four monkeys (B, F, I, and S) performed the task and completed 33/1166, 30/1080, 30/1080, and 28/1008 sessions and blocks, respectively. We used a token-based rewarded multi-dimensional attention task in which monkeys were required to explore the objects on the screen and determine through trial and error a target feature while learning to ignore irrelevant dimensions and features. Stimuli were multi-dimensional 2D viewed Quaddles which could present 1 to 3 features other than a neutral Quaddle (**Fig. 1C**) (Watson, Voloh, Naghizadeh, et al., 2019). Four different dimensions/features were used in the task design (Shape, Color, Arm, and Pattern) and in each session of learning features of three dimensions were selected as potential target and distractor features for the learning blocks of that session. In each learning block only one feature corresponded to the correct target. All objects had the same number of non-neutral features. Attentional load was indexed as the number of non-neutral features presented on each object, varying from 1 to 3. The targeted feature was un-cued and monkeys were required to find the targeted feature through trial and error in each learning block. The task rule (target feature) remained the same throughout a learning block. After each block of learning the task rule changed and monkeys had to explore again to find out the new rule.

Each correct touch was followed by a yellow halo around the stimulus as visual feedback (500 ms), an auditory tone, and a fixed number of animated tokens traveling from the chosen object location to the token bar on top of the screen (**Fig. 1A**). Erroneously touching a distractor object was followed by a blue halo around the touched objects, a low-pitched auditory feedback, and in the loss conditions travelling of one or three empty (grey) tokens to the token bar where the number of lost tokens were removed from the already earned tokens as an error penalty (feedback timing was the same as for correct trials). To receive a fluid reward, i.e. to cash out the tokens, monkeys had to complete 5 tokens in the token bar. When five tokens were collected the token bar blinked red/blue three times, a high pitch tone was played as auditory feedback and fluid was delivered. After cashing out the token bar was reset to five empty token placeholders.

### Experimental design

In each learning block one fixed token reward schedule was used, randomly drawn from seven distinct schedules (see below). We simulated the token schedules to reach a nearly evenly spaced difference in reward rate while not confounding the number of positive or negative tokens with reward rate (**Fig. 1D,E**). For simulating the reward rate with different token gains and losses we used a tangent hyperbolic function to simulate a typical learning curve by varying the number of trials needed to reach 75% accurate performance criterion from 5 to 25 trials (the so called ‘learning point’ in a block). This is the range of learning points the subjects showed for the different attentional load conditions in 90% of blocks during training. We simulated 1000 blocks of learning for different combinations of gained- and lost-tokens for correct and erroneous choices, respectively. For each simulated token combination, the reward rate was calculated by dividing the frequency of the full token bar to the block length. The average reward rate was then computed as the average reward rate over all simulation runs for each token condition. The reward rate showed on average what is the probability of receiving reward per trial. Seven token conditions were designed with a combination of varying gain (G; 1, 2, 3, and 5) and loss (L; 3, 1, and 0) conditions (1G 0L, 2G - 3L, 2G -1L, 2G 0L, 3G -1L, 3G 0L, and 5G 0L). The 5G/0L condition entailed providing 5 tokens for a correct choice and no tokens lost for incorrect choices. This condition was used to provide animals with more opportunity to earn fluid reward than would be possible with the conditions that have lower reward rate. Since gaining 5 tokens were immediately cashed out for fluid delivery, we do not consider this condition for the token-gain and token-loss analysis as it confounds secondary and primary reinforcement in single trials.

### Analysis of learning

The improvement of accuracy over successive trials relative to the beginning of a learning block reflects the learning curve, which we computed with a forward-looking 10-trial averaging window in each block (**Fig. 2A,C,E**). We defined learning in a block as the number of trials needed to reach criterion accuracy of ≥75% correct trials over 10 successive trials. Monkeys on average reached learning criterion, i.e. learnt, >75% of blocks (B=67%, F=75%, I=83%, and S=74%, *see* **Supplementary Fig. 3**). We computed a *post-learning accuracy* as the proportion of correct choices in all trials after the learning criterion was reached in a block (**Fig. 4**).

**Figure 2.**
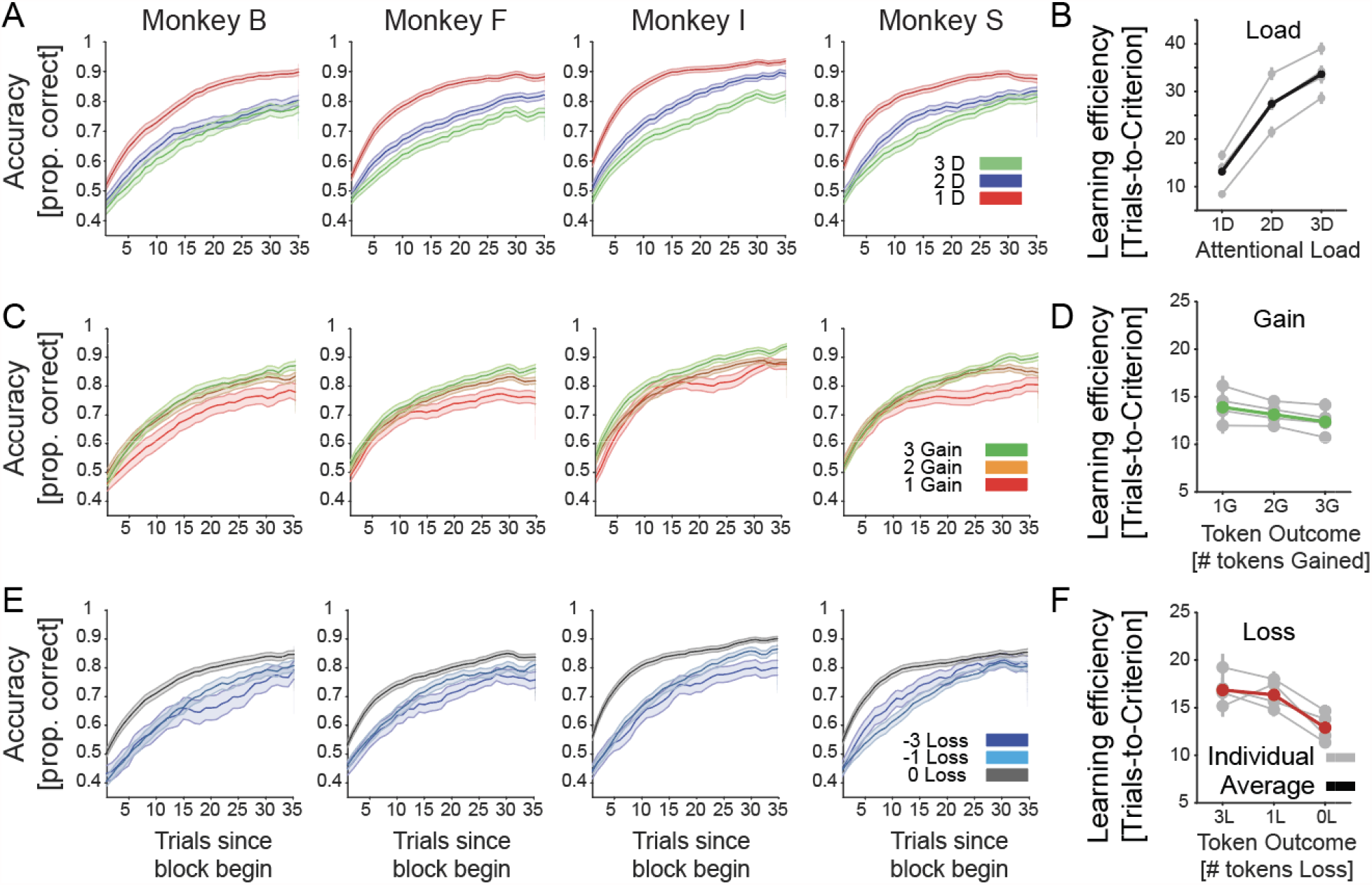
Average learning curves for each monkey and load, loss, and gain conditions. **A:** The proportion correct performance (*y-axis*) for low/medium/high attentional load for each of four monkeys relative to the trial since the beginning of a learning block (*x-axis*). **B**: The number of trials-to-reach criterion (*y-axis*) for low/medium/high attentional load (*x-axis*) for each monkey (in *grey*) and their average (in *black*). **C:** Same as A showing the learning for blocks in which 1, 2, or 3 tokens were gained for correct performance. Red line shows average across monkeys. **D:** Same as *B* for blocks where monkeys expected to lose 0, 1, or 3 tokens for incorrect choices. **E:** Same as A and B showing the learning for blocks in which 0, 1, or 3 tokens were lost for incor ect performance. Green line shows average across monkeys. Errors are SE’s. **F:** Same as B blocks where monkeys expected to win 1, 2, or 3 tokens for correct choices.

**Figure 3.**
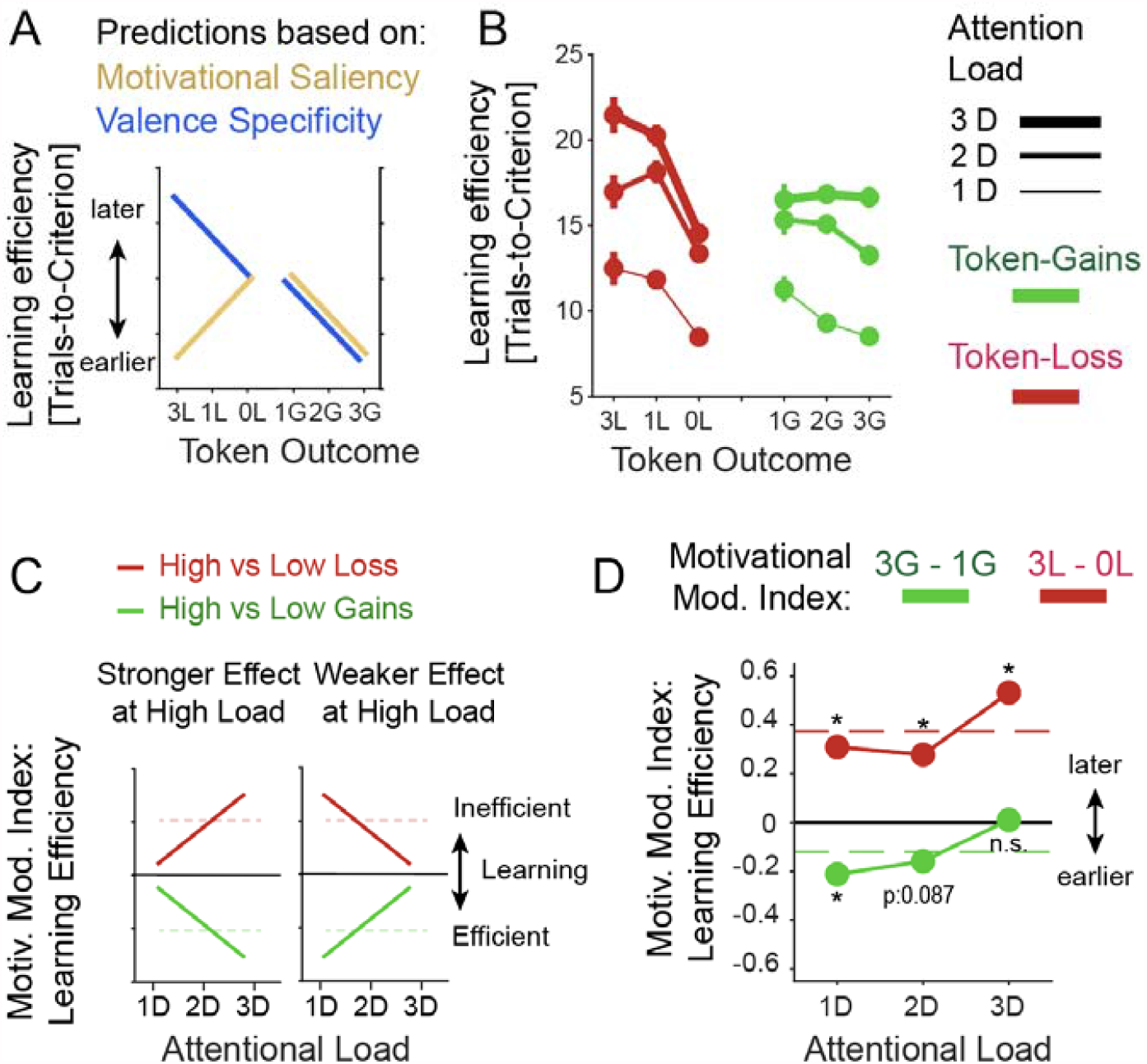
The effect of attentional load and expected token gain/loss on learning efficacy. **A:** The motivational saliency hypothesis (*orange*) predicts that learning efficacy is high (learning criterion is reached early) when the absolute magnitude of expected loss (to the left) and expected gains (to the right) is high. The valence-specific hypothesis (*blue*) predicts that learning efficacy is improved with high expected gains (to the right) and decreased with larger penalties/losses. **B**: Average learning efficacy across four monkeys at low/medium/high attentional load (line thickness) in blocks with increased expected token-loss (to the left, in red) and with increased expected token gains (to the right, green). **C:** Hypothetical interactions of expected gains/losses and attentional load. Larger incentives/penalties might have a stronger positive/negative effect at higher load (*left*) or a weaker effect at higher load (*right*). The predictions can be quantified with the Motivational Modulation index, which is the difference of learning efficacy for the high vs. low gains conditions (or high vs low loss conditions). **D:** Average motivational modulation index shows that the slowing effect of larger penalties increased with higher attentional load (*red*). In contrast, the enhanced learning efficacy with higher gain expectations are larger at lower attentional load and absent at high attentional load (*green*).

**Figure 4.**
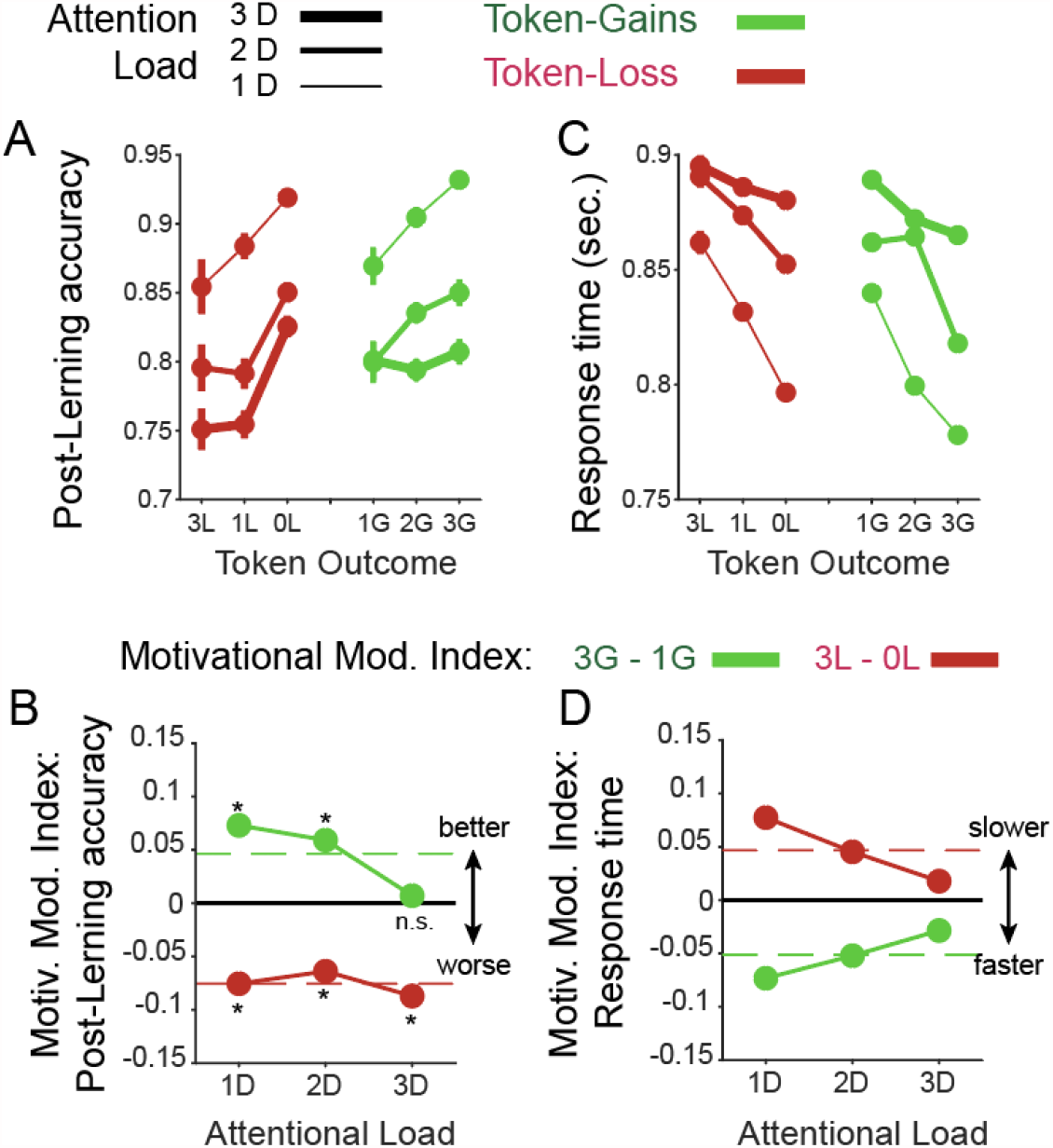
The effect of attentional load and token gain/loss expectancy on post-learning performance and response times. A: Post-learning accuracy (*y-axis*) when expecting varying token loss (*red*) and gains (*green*) at low/medium/high attentional load (line thickness). Overall, learning efficiency decreased with larger penalties and improved with larger expected token-gains. B: The motivation modulation index for low/medium/high attentional load (*x-axis*) shows that the improvement with higher gains was absent at high load, and the detrimental effect of penalties on performance was evident at all loads. C,D: Same format as A,B for response times. Subject slowed down when expecting larger penalties and speed up responses when expecting larger gains (C). These effects were largest at low attentional load and decreased at higher load (D).

### Statistical analysis

We first constructed a linear mixed effect model (Pinherio & Bates, 1996) that tests how learning speed (indexed as the ‘learning trial’ (LT) at which criterion performance was reached) and accuracy after learning (*LT/Accuracy*) over blocks are affected by 3 factors, *attentional load (Att*_*Load*_) with three levels (1D, 2D, and 3D distractor feature dimensions), the factor feedback gain (*Fb*_*Gain*_) with 3 levels (Gaining tokens for correct performance: 1, 2, or 3), and the factor feedback loss (*Fb*_*Loss*_) with 3 levels (Loss of tokens for erroneous performance: 0, -1, or -3). We additionally considered as random effects the factor *Monkeys* with 4 levels (B, F, I, and S), and the factor *target features* (*Feat*) with 4 levels (color, pattern, arm, and shape). The random effects control for individual variations of learning and for possible biases in learning features of some dimensions better or worse than for others. This LME had the form

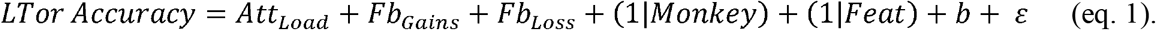

The model showed significant main effects of all three factors and random effects laid inside the 95% confidence interval (**Supplementary Fig. 3**). To test for the interplay of motivational variables and attentional load we extended the model with the interaction effects for *Att*_*Load*_ x *Fb*_*Gain*_ and *Att*_*Load*_ x *Fb*_*Loss*._ to:

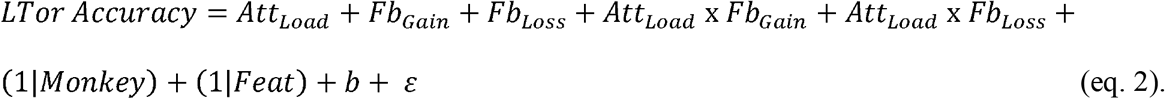

To compare the model with and without interaction terms we used Bayesian information criterion (BIC) as well as the Theoretical Likelihood Ratio Test (Hox, 2002) to decide which model explains the data best.

We also tested two additional models that tested whether the absolute difference of *Fb*_*Gain*_ and *Fb*_*Loss*_ played a significant role in accounting for accuracy and learning speed using (*Diff*_*Gain-Loss*_ = *Fb*_*Gain*_ - *Fb*_*Loss*_) as predictor. Secondly, we tested the role of Reward Rate (*RR*) as predictor. However, both these two models were inferior to the LME’s formulated in eq. 2 and showed lower BIC. We thus do not describe these factors further.

In addition to the block-level analysis of learning and accuracy (eq. 1-2), we also investigated the trial-level to quantify how accuracy and the response times (*Accuracy/RT*) of the monkeys over trials are modulated by four factors. The first factor was the learning status (*Learn*_*State*_) with two levels (before and after reaching the learning criterion). In addition, we used the factor *attentional load (Att*_*Load*_) with three levels (1D, 2D, and 3D objects), the factor feedback gain (*Fb*_*Gain*_) on the previous trial with three levels (Gaining tokens for correct performance: 1/2/3), and the factor feedback loss on the previous trial (*Fb*_*Loss*_) with three levels (Loss of tokens for erroneous performance: 0/-1/-3).

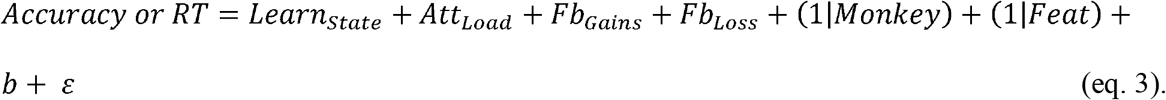

Eq. 3 was also further expanded to account for interaction terms *Att*_*Load*_ x *Fb*_*Gain*_ and *Att*_*Load*_ x *Fb*_*Load*_.

In order to quantify the effectiveness of the token bar to modulate reward expectancy we predicted accuracy or RT of the animals by how many tokens were already earned and visible in the token bar, using the factor *Token* _*state*_ defined as the number tokens visible in the token-bar as formalized in eq. 4 and shown in **Fig. 4**.

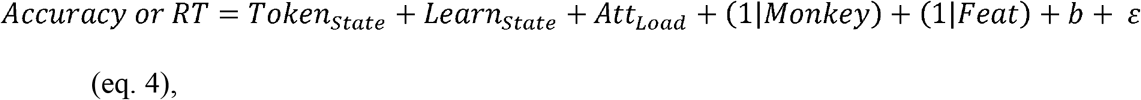

We compared two models with and without including the factor *Token*_*State*_. We then ran 500 simulations of likelihood ratio tests and found that the alternative model which included the factor *Token*_*State*_ had a better performance than the one without the factor *Token*_*State*_ (p<0.002, LRstat=1100.7, BIC_without Token_state_ =19840, BIC_without Token_state_ =19949).

### Analyzing the motivational modulation of attentional load

To evaluate the influence of increasing gains and increasing losses on attentional load we calculated Motivational Modulation Indices (MMI) as the difference in learning speed (the trial-to-criterion) in the conditions with 3 versus 1 token gains (MMI_Gain_) or in conditions with 3 versus 0 token losses (MMI_Loss_). MMI_Gain_ and MMI_Loss_ were calculated separately for the low, medium and high attentional load condition. We statistically tested these indices under the null hypothesis that they are not different from zero by performing pairwise t-tests on each gain or loss condition given low, medium, and high attentional load (1D, 2D, and 3D).

To validate the t-test MMI results at individual load conditions, we used a permutation approach that tested the MMI across attentional load conditions. At the first step we randomly selected 100 samples from each feedback gain/loss condition and computed the MMI for each pair of samples separately for the feedback gain and for the feedback loss conditions in each attentional load. For each attention load we selected 1000 random sub-samples with a sample size of 100 from each of the other 2 attentional loads separately, constructed 1000 95% confidence intervals and chose the top 5% confidence intervals to rectify the multiple comparison correction under the null hypothesis that MMI in each attentional load condition is not different from the each of other attentional load conditions.

## Results

Four monkeys performed the feature learning task and collected tokens as secondary reinforcers to be cashed out for fluid reward when five tokens were collected. The task required learning a target feature in blocks of 35-60 trials through trial-and-error by choosing one among three objects composed of multiple different object features (**Fig. 1A,B**). Attentional load was varied by increasing the number of distracting features of these objects to be either only from the same feature dimension as the rewarded target feature (‘*1-Dimensional*’ load), or additionally from a second feature dimension (‘*2-Dimensional*’ load), or from a second and third feature dimension (‘*3-Dimensional’* load). We achieved this parametric variation of distractibility using multidimensional Quaddle objects that varied features of 1, 2, or 3 dimensions from trial-to-trial (**Fig. 1C**)(Watson, Voloh, Naghizadeh, et al., 2019). In addition to load, we orthogonally varied between blocks the number of tokens that were gained for correct responses (1, 2, or 3 tokens), or that could be lost for erroneous responses (0, -1, -3 tokens). We selected combinations of gains and losses so that losses were used in a condition with relatively high reward rate (e.g. a condition with 3 gain and 1 loss tokens), while other conditions had lower reward rate despite the absence of loss tokens (e.g. condition with 1 gain and 0 loss tokens). This arrangement allowed studying losses relatively independent of overall reward rate (**Fig. 1D,E**).

During learning monkeys showed slower choice response times the more tokens were already earned. This suggests that they tracked the tokens they obtained and were more careful responding the more they had earned in those trials in which they were not yet certain about the rewarded feature (**Supplementary Fig. S1A,C**). After they reached the learning criterion (during asymptotic performance) monkeys showed faster response times the more tokens they had earned (**Supplementary Fig. S1B,D**, LME models including the variable *“ Token*_*State*_ *”* explained the response time better than those without *“ Token*_*State*_ *”* variable, p<0.002, LR_stat_=1100.7, BIC_without Token_state_=19840, BIC_without Token_state_=19949). On average each subject completed 1080 learning blocks (SE ±32, range 1008-1166) in 30 test sessions (SE ±1, range 28-33). All monkeys showed slower learning of the target feature with increased attentional load and when experiencing losing more tokens for incorrect responses, and all monkeys showed increased speed of learning the target feature the more tokens they could earn for correct responses (**Fig. 2**; the same result pattern was evident for reaction times, **Supplementary Fig. S2)**. The effects of load, loss, and gains were evident in significant main effects of linear mixed effects models (LME’s, see Methods, Att._Load_: b= 4.37, p<0.001; feedback_Losses_: b=1.39, p=<0.001; feedback_Gains_: b=-0.76, p=0.008). In all linear mixed effects models monkeys (numbered 1-4) and the target feature dimensions (arms, body shapes, color, and surface pattern of objects) served as random grouping effects. No significant random effects were observed unless explicitly mentioned. We interpret the main effects of learning the target feature at varying distractor load and gains/losses as reflecting changes in the learning efficacy. Not only did monkeys learn faster at lower loads, when expecting less losses, and when expecting higher gains, but they also had fewer unlearned blocks under these same conditions (**Supplementary Fig. 3**).

The main effects of prospective losses and gains provide apparent support for a valence-specific effect of motivational incentives and penalties on attentional efficacy. They are not easily reconciled with the ‘loss attention’ framework because increasing losses impared rather than enhanced learning (**Fig. 3A**) (Yechiam & Hochman, 2013b; Yechiam, Retzer, Telpaz, & Hochman, 2015). However, the valence-specific effect was not equally strong at low/medium/high attentional loads (**Fig. 3B**). We found that linear mixed effects (LEM) models with interaction terms described the data better than without them (Likelihood Ratio Stat. = 8.5, p=0.014, Bayesian Inform Crit. 21176 and 21169 for [Att._Load_ x (feedback_Losses_ + feedback_gains_)], and [Att._Load_ + feedback_Losses_ + feedback_gains_], respectively). To visualize these interactions, we calculated the Motivational Modulation Index (MMI) as the difference in learning efficacy (avg. number of trials-to-criterion) when expecting to gain 3 vs. 1 tokens for correct choices [MMI_Gains_ = LEfficacy_3G_-LEfficacy_1G_], and when experiencing losing 3 vs 0 tokens for incorrect choices [MMI_Loss_ = LEfficacy_3L_-LEfficacy_0L_]. By calculating the MMI for each attentional load condition we can visualize whether the motivation effect of increased prospective gains and losses increases or decreases with higher attentional load (**Fig. 3C**). We found that the detrimental effect of larger prospective losses on learning increased with attentional load, causing a larger decrease in learning efficacy at high load (**Fig. 3D**) (permutation test, p < 0.05). In contrast, expecting higher gains improved learning most at low attentional load and had no measurable effect at high load (**Fig. 3D**) (permutation test, p < 0.05). Pairwise t-test comparison also showed MMI_Loss_ was significantly different from zero (p=0.002, p=0.008, and p=0.001 for low, medium, and high load), while MMI_Gains_ was only significantly different from zero in the low load gain condition (p=0.007, p=0.087, and p=0.95 for low, medium, and high load).

These contrasting effects of gains and losses on learning efficacy were partly paralleled in post-learning accuracy. Accuracy was enhanced with higher expected gains at lower but not at the highest attentional load conditions (t-test pairwise comparison, p<0.001, p=0.01, and p=0.78 for low, medium, and high load), decreased with higher expected losses at all loads (t-test pairwise comparison, p=0.003, p=0.017, and p=0.001 for low, medium, and high load), but this decrease was not modulated by load level (permutation test, p>0.05) (**Fig. 4A,B**). In contrast to learning speed and post-learning accuracy, response times varied more symmetrically across load conditions. At low load choice times were fastest with larger expected gains, and slowest with larger expected token-loss (**Fig. 4C,D**). At medium and higher attentional loads these effects were less pronounced.

To understand how prospective gains and losses modulated learning efficacy on a trial-by-trial level we calculated the choice accuracy in the trials immediately after experiencing a loss of 3, 1, or 0 tokens, and after experiencing a gain of 1, 2, or 3 tokens. Experiencing more negative outcomes caused all monkeys to improve performance in the trials immediately following the negative outcome (**Fig. 5A**). This finding shows subjects improved performance after losing tokens. The behavioral improvement after having lost tokens was particularly apparent in the low attentional load condition, and less in the medium and high attentional load conditions (**Fig. 5B**) (LME model predicting the previous trial effect on accuracy; [Att._Load_ x feedback_Losses_], b=0.01, p=0.002). We quantified this effect by taking the difference in performance for loosing 3 vs. 0 tokens, which confirmed that there was maximal benefit from larger penalties in the low load condition and least influence in the high load condition (**Fig. 5C**). Similar to losses, experiencing larger gains improved the accuracy in the subsequent trial strongest at low and medium attentional load (LME model predicting previous trial effects on accuracy: for [Att._Load_ x feedback_Gains_], b=-0.03, p<0.001) (**Fig. 5B,C**). Thus, motivational improvements of post-token feedback performance adjustment was evident for token gains as well as for losses but primarily evident at lower and not at higher attentional load.

**Figure 5.**
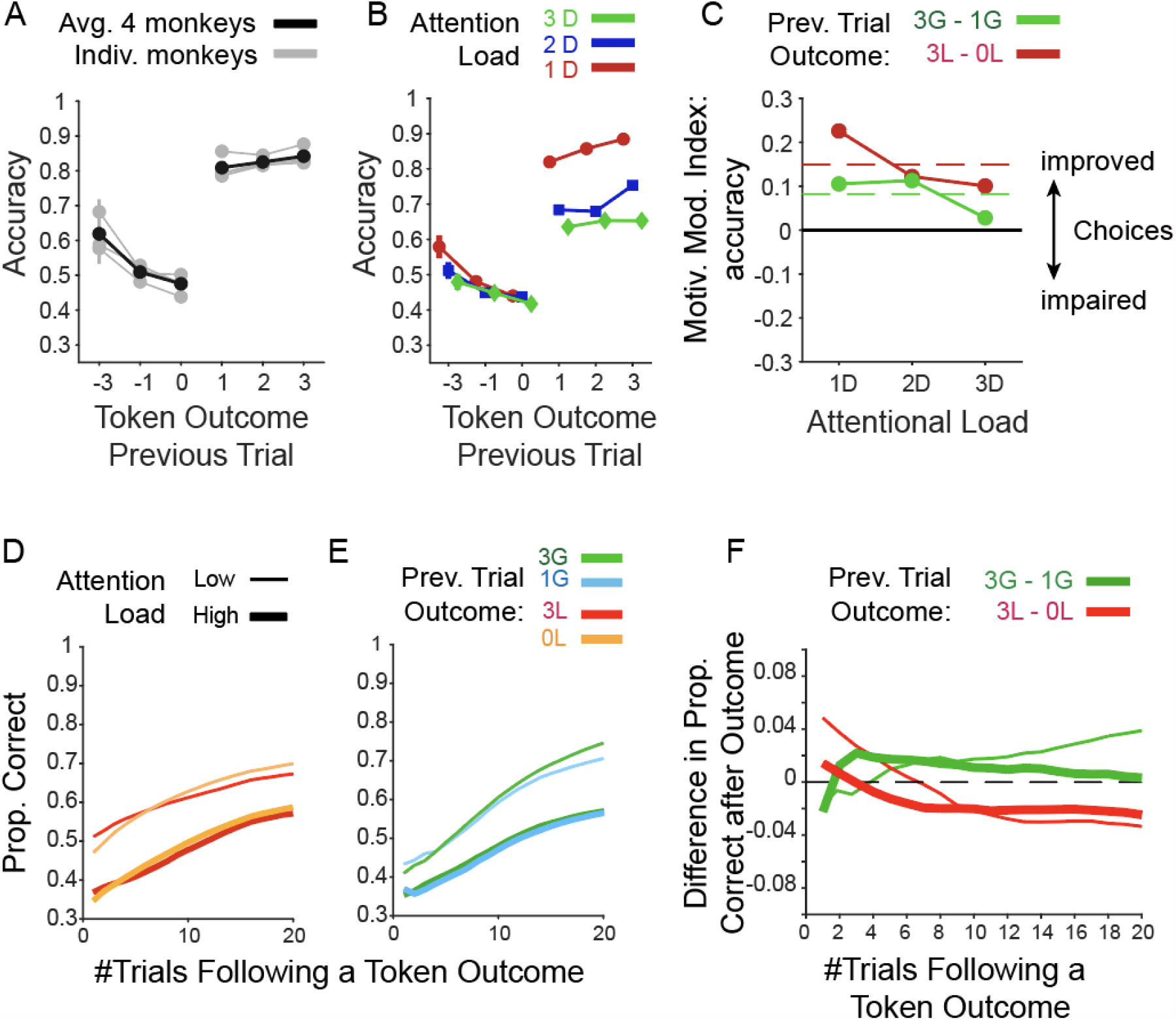
Immediate effects of experienced token gains and losses on performance. **A**: The effects of an experienced loss of 3, 1 or 0 tokens and of an experienced gain of 1, 2, or 3 tokens (*x-axis*) on the subsequent trials’ accuracy (*y-axis*). Grey lines are from individual monkeys, and black shows their average. **B**: Same format as *A* for the average previous trial outcome effects for low/medium/high attentional load. **C:** Motivational modulation index (y-axis) quantifies the improved accuracy after experiencing 3 vs. 1 losses (*red*), and after experiencing 3 vs. 1 token gains (*green*) for low/medium/high attentional load. **D**: The effect of an experienced loss of -3 or 0 token during learning on the proportion of correct choices in the following n^th^ trials (x-axis). **E:** Same as *D* but for an experienced gain of 3 or 1 token. **F**: The difference in accuracy (*y-axis*) after experiencing 3 vs. 0 losses (*red*), and 3 vs. 1 token gains (*green*) over n trials subsequent to the outcome (x-axis). Thick and thin lines denote low and high attentional load conditions, respectively.

So far, the previous trial analysis considered the accuracy only in the immediate trial following lost or gained tokens. To discern longer term effects, we calculated accuracy with a n-trial running average window for n = 1-20 trials following the token outcome during learning (trials before the learning criterion is reached). The analysis showed that immediately after losing 3 tokens, accuracy was increased compared with no losses, however this effect was transient and reversed within 2 trials in the high load condition, and within 6 trials in the low load conditions (**Fig. 5D,F**). For the gain-conditions the results were different. A larger gain of 3 (vs. 1) token led to longer-lasting improvement of performance. This improvement was longer lasting in the low than the high attentional load condition (**Fig. 5E-F**). These longer-lasting effects on performance closely resemble the main effects of losses and gains on the learning efficacy (**Fig. 2, 3**) and suggest that the block level learning effects be traced back to outcome-triggered performance changes at the single trial level with stronger negative effects after larger losses and stronger positive effects after larger gains.

## Discussion

We found that prospective gains and losses had opposite effects on learning a relevant target feature. Experiencing losing tokens reduced learning speed (trials to criterion), impaired retention (post-learning accuracy), and increased choice response times, while experiencing gaining tokens had the opposite effects. These effects varied with attentional load in opposite ways. Larger penalties for incorrect choices had maximally detrimental effects when there were many distracting features (high load). Conversely, higher gains for correct responses enhanced flexible learning at lower attentional load but had no beneficial effects at higher load. These findings were paralleled on the trial level, which showed reduced behavioral adjustment following errors when losing 3 compared to 1 token, and by a lack of improving performance after correct choices of targets in the high load condition.

Together, these results document apparent asymmetries of gains and losses on learning efficacy. The negative effects of losing tokens and the positive effects of gaining tokens were apparent in all four monkeys. The inter-subject consistency of the token and load effects suggests that our results delineate a fundamental relationship between the motivational influences of incentives (gains) and disincentives (losses) on learning efficacy at varying attentional loads.

### Experiencing loss reduces learning efficacy

One key observation in our study is that rhesus monkeys are sensitive to visual tokens as secondary reinforcers, closely tracking their already obtained token assets (**Supplementary Fig. 1**). This finding is consistent with prior studies in rhesus monkeys showing that gaining tokens for correct performance and losing tokens for incorrect performance modulates performance in choice tasks (Rich & Wallis, 2017; Seo & Lee, 2009; Taswell et al., 2018). This sensitivity was essential to functionally separate the influence of negative and positive expectancies from the influence of the overall saliency of outcomes. In our study the number of tokens for correct and incorrect performance remained constant within blocks of ≥35 trials and thus set a reward context for learning target features among distractors in our task (Sallet et al., 2007). When this reward context entailed losing three tokens, monkeys learned the rewarded target feature ∼5-7 trials later (slower) than when incorrect choices led to no token change (depending on load, see **Fig. 3B**). This finding shows that experiencing losing 3 tokens does not enhance the processing, or the remembering, of the chosen distractors, but rather impairs using information from the erroneously chosen objects to adjust behavior. This finding had a relatively large effect size and was rather unexpected given various previous findings that would have predicted the opposite effect. First, the *loss attention* framework suggests that experiencing a loss causes an automatic vigilance response that triggers subjects to explore alternative options other than the chosen object (Yechiam et al., 2019; Yechiam & Hochman, 2013a, 2013b). Such an enhanced exploration might have facilitated avoiding objects with features that were associated with the loss. But our results suggest that the assumed loss-triggered re-orienting to alternative objects was either not enhanced after loosing tokens, or that loosing tokens impaired encoding the specific object features of the loss-inducing object so that the animals were less able to distinguish which objects in future trials belonged to the loss-inducing objects. This poorer encoding would lead to less informed exploration after a loss, which would not facilitate but impair learning. This account is consistent with human studies showing poorer discrimination of stimuli when they are associated with monetary loss, aversive images or odors, or electric shock (Laufer et al., 2016; Laufer & Paz, 2012; Resnik et al., 2011; Schechtman et al., 2010; Shalev et al., 2018).

A second reason why the performance decrement with loss was unexpected is because monkeys and humans clearly show non-zero (>0) learning rates for negative outcomes when they are estimated separately from learning rates for positive outcomes (Caze & van der Meer, 2013; Collins & Frank, 2014; Frank, Moustafa, Haughey, Curran, & Hutchison, 2007; Frank, Seeberger, & O’Reilly R, 2004; Seo & Lee, 2009), indicating that negative outcomes in principle are useful for improving the updating of value expectations. While we found that subjects did improve performance immediately after an incorrect performance, this improvement disappeared after ∼3 further trials and overall caused slower learning. One possibility is that our task reduced not the learning rates per se, but the credit assignment process that updates the expected values of object features based on negative or positive outcomes. This suggestion calls upon future investigations to clarify the nature of the loss-induced impediment using computational modeling of the specific subcomponent processes underlying the learning process (Womelsdorf, Watson, & Tiesinga, 2021).

A third reason why loss-induced impairments of learning were unexpected are prior reports that monkeys successfully learn to avoid looking at objects that are paired with negative reinforcers (such as a bitter taste) (Ghazizadeh et al., 2016b). According to this prior finding, monkeys should have effectively avoided choosing objects with loss-inducing features when encountering them again. Instead, the token loss reduced their capability to avoid the objects sharing features with the object that caused token loss in previous trials. The discrepancy with our findings might again lie in the complexity of the objects used in our task that made it difficult to encode and memorize the precise features of the chosen loss-inducing objects.

The three discussed putative reasons for why loss might not have improved but decreased performance point to the complexity of the object space we used. When subjects lost already attained tokens for erroneously choosing an object with 1, 2, or 3 object features, they were less able to assign negative credit to a specific feature of the chosen object. Consequently, they did not learn from erroneous choices as efficiently as they would have learned with no or neutral feedback after errors. This account is consistent with studies showing a wider generalization of aversive outcomes and a concomitant reduced sensitivity to the loss-inducing stimuli (Laufer et al., 2016; Laufer & Paz, 2012; Resnik et al., 2011; Schechtman et al., 2010; Shalev et al., 2018). Consistent with such reduced encoding or impaired credit assignment of the specific object features, we found variable effects on post-error adjustment (**Fig. 5B,C**) and reduced longer-term (i.e. over ∼5-20 trials) performance after experiencing loss (**Fig. 5F**) with the negative effects increasing with more distractors in the higher attentional load condition (**Fig. 3**). According to this interpretation, penalties such as losing tokens are detrimental to performance when they do not inform straightforwardly what type of features should be avoided. This view is supported by additional evidence. Firstly, when tasks have simpler, binary choice options, negative outcomes are more immediately informative about which objects should be avoided and learning from negative outcomes can be more rapid than learning from positive outcomes (Averbeck, 2017). We found a similar improvement of performance in the low load condition that lasted for 1-3 trials before performance declined below average (**Fig. 5F**). Secondly, using aversive outcomes in a feature non-selective way might incur a survival advantage in various evolutionary meaningful settings. When a negative outcome promotes generalizing from specific aversive cues (e.g. encountering a specific predator) to a larger family of aversive events (all predator-like creatures) this can enhance fast avoidance responses in future encounters (Barbaro et al., 2017; Dunsmoor & Paz, 2015; Krypotos, Effting, Kindt, & Beckers, 2015; Laufer et al., 2016). Such a generalized response is also reminiscent of non-selective ‘escape’ responses that experimental animals show early during aversive learning before they transition to more cue specific avoidance responses later in learning (Maia, 2010). The outlined reasoning helps explaining why we found that experiencing loss is not helpful to avoid objects or object features when multiple, multidimensional objects define the learning environment.

### Experiencing gains enhances attentional efficacy but can’t compensate for distractor overload

We found that experiencing three tokens as opposed to one token for correct choices improved the efficacy of learning relevant target features by ∼4, ∼1.5, and ∼0 trials in the low, medium and high attentional load condition (**Fig. 3**). On the one hand, this finding provides further quantitative support that incentives can improve learning efficacy (Berridge & Robinson, 2016; Ghazizadeh et al., 2016b; Walton & Bouret, 2019). In fact, across conditions learning was most efficient when the monkeys expected three tokens and when objects varied in only one feature dimension (1D, low attentional load condition). However, this effect disappeared at high attentional load, i.e. when objects varied trial-by-trial in features of three different dimensions (**Fig. 3**). A reduced behavioral efficiency of incentives in light of distracting information is a known phenomenon in the problem solving field (Pink, 2009), but an unexpected finding in our task because the complexity of the actual reward rule (the rule was ‘*find the feature that predicts token gains*’) did not vary from low to high attentional load. The only difference between these conditions was the higher load of distracting features, suggesting that the monkeys might have reached a limitation in distractor interference that they could not compensate by mobilizing additional control resources.

But what is the source of this limitation to control distractor interference? One possibility is that when attention demands increased in our task monkeys are limited in allocating sufficient control or ‘mental effort’ to overcome distractor interference (Hosking, Cocker, & Winstanley, 2014; Klein-Flugge, Kennerley, Friston, & Bestmann, 2016; Parvizi, Rangarajan, Shirer, Desai, & Greicius, 2013; Rudebeck, Walton, Smyth, Bannerman, & Rushworth, 2006; Shenhav et al., 2017; Walton, Bannerman, Alterescu, & Rushworth, 2003; Walton, Rudebeck, Bannerman, & Rushworth, 2007). Thus, subjects might perform poorer at high attentional load partly because they are not allocating sufficient control on the task (Botvinick & Braver, 2015). Rational theories of effort and control suggest that subjects allocate control as long as the benefits of doing so outweigh the costs of exerting more control and perform poorer at higher demand if the benefits (e.g. the reward rate for correct performance) are not increased concomitantly (Shenhav, Botvinick, & Cohen, 2013; Shenhav et al., 2017). The strength of such a rational theory of attentional control is that it does propose a specific, biologically plausible feature that limits control, which is assumed to be the degree of cross-talk of conflicting information at high cognitive load (Shenhav et al., 2017). According to this hypothesis, effort and control demand corresponds to the degree of interfering information. This view provides a parsimonious explanation for our findings at high attentional load. While incentives can increase control-allocation when there is little cross-talk of the target feature with distractor features (at low attentional load), the incentives are not able to compensate for the increased cross-talk of distracting features at high attentional load condition. Our results therefore provide quantitative support for a rational theory of attentional control. In our task the critical transition from sufficient to insufficient control of interference was between the medium and high attentional load condition, which corresponded to an increase of distractor features that vary trial-by-trial from 8 features (at medium attentional load) to 26 features (at high attentional load). Thus, subjects were not able to compensate for distractor interference when the number of interfering features were between 8 and 26, suggesting that prospective gains – at least when using tokens as reinforcement currency - hit a limit to enhance attentional control within this range.

### Impaired Learning in loss-contexts and economic decision theory

In economic decision theory it is well documented that making choices are suboptimal in contexts that frame outcomes in terms of losses rather than gains (Tversky & Kahneman, 1981, 1991). This view from behavioral economics aims to explain which options individuals will choose in a given context, which shares some resemblance with our task, where subjects learned to choose one among three objects in learning contexts (blocks) that were framed with variable token gains and losses. In a context with higher potential losses, humans are less likely to choose an equally valued option than in a gain-context. The reason for this irrational change in choice preferences is believed to reside in an overweighting of emotional content that devalues outcomes in loss contexts (Barbaro et al., 2017; Loewenstein, Weber, Hsee, & Welch, 2001). Colloquial words for such emotional overweighting might be displeasure (Tversky & Kahneman, 1981), distress, annoyance, or frustration. Concepts behind these words may relate to primary affective responses of anger, disgust or fear. However, these more qualitative descriptors are not providing an explanatory mechanism, but rather tend to anthropomorphize the observed deterioration of learning in loss-contexts. Moreover, the economic view does not provide an explanation why the learning would be stronger affected at higher attentional load (**Fig. 3D**).

## Conclusion

Taken together, our results document the interplay of motivational variables and attentional load during flexible learning. We first showed that learning efficacy is reduced when attentional load is increased despite the fact that the complexity of the feature-reward target rule did not change. This finding illustrates that cognitive control processes cannot fully compensate for an increase in distractors. The failure to fully compensate for enhanced distraction was not absolute. Incentive motivation was able to enhance learning efficacy when there were distracting features of one or two feature dimensions but could not help anymore to compensate for enhanced interference when features of a third feature dimension intruded into the learning of feature values. This limitation suggests that crosstalk of distracting features represents a key process involved in cognitive effort (Shenhav et al., 2017). Moreover, the negative effect of distractor interference was exacerbated by experiencing the loss of tokens for wrong choices. This effect illustrates that negative feedback does not help to avoid loss-inducing distractor objects during learning, which docunents that expecting or anticipating loss deteriorates flexible learning the relevance of objects in a multidimensional object space.

## Acknoweldgements

The authors would like to thank Adam Neuman and Seyed-Alireza Hassani for help collecting the data. Research reported in this publication was supported by the National Institute of Mental Health of the National Institutes of Health under Award Number R01MH123687. The content is solely the responsibility of the authors and does not necessarily represent the official views of the National Institutes of Health.

## Author Contributions

K.B.B. and T.W. conceived and performed the experiments. M.R.W., T.W. and K.B.B. wrote the code to run the experiments. K.B.B. analyzed the data. T.W., K.B.B., and M.R.W. wrote the paper.

## Data and code accessibility

All data supporting this study and its findings, as well as custom MATLAB code generated for analyses, are available from the corresponding author upon reasonable request.

## Supplementary Information

### Supplementary Figures S1-S3

**Supplementary Figure 1.**
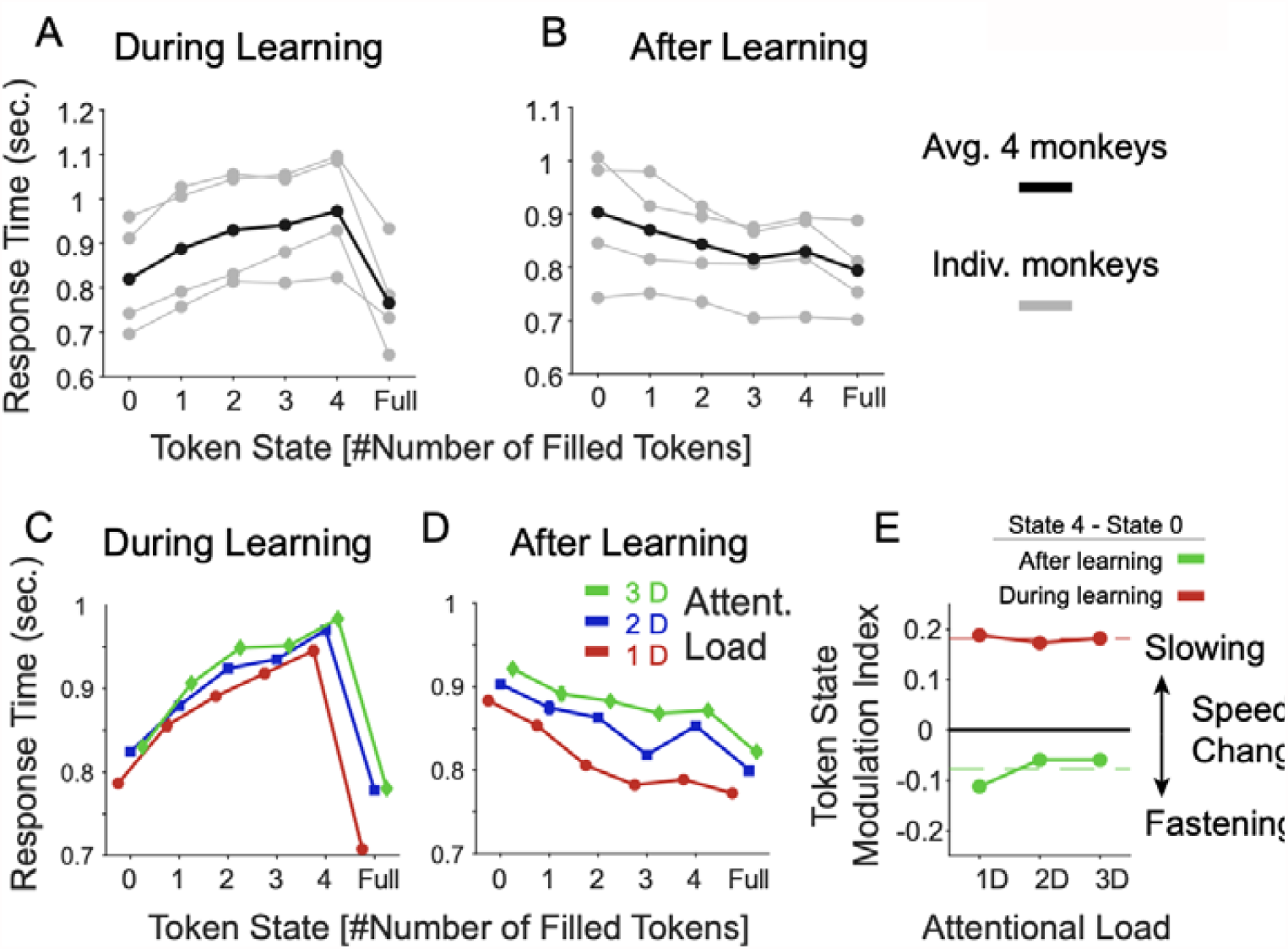
Effects of the number of earned tokens (‘Token State’) on response times. **A**: Average response times as a function of the number of earned tokens visible in the token bar (*x-axis*) for individual monkeys (grey) and their average (in black). Included were only trials prior to reaching the learning criterion. Over all trials monkeys (B, F, I, and S) showed average response time of 785, 914, 938, and 733 ms, respectively. **B**: Same as *A* but including only trials after the learning criterion was reached. **C-D:** Same format as *A-B* showing the average response times across monkeys for the low, medium, and high attentional load condition during learning (*C*) and after learning (D). **E:** The Token State modulation index (*y-axis*) shows the difference in response times (RT’s) when the animal had 4 tokens earned versus one token earned. During learning (*red*) RT’s were slower with 4 than 1 earned tokens to similar degrees for different attentional loads (x-axis). This pattern reversed after learning was achieved (*green*).

**Supplementary Figure 2.**
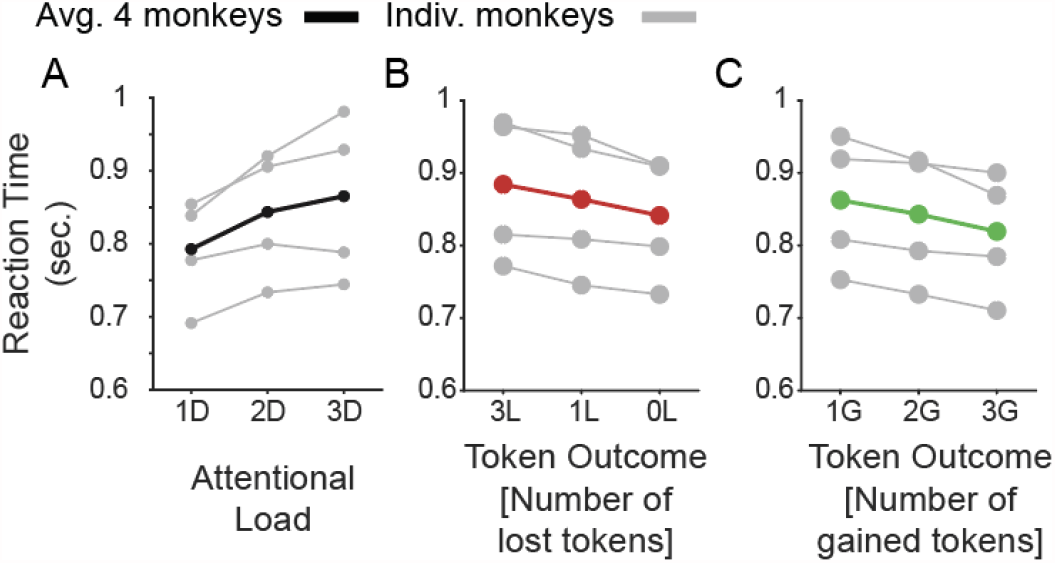
Main effects of attentional load, number of expected token-gains and expected token-loss on learning speed and response time. **A**: The response times (*y-axis*) for low/medium/high attentional load (*x-axis*) for each monkey (in *grey*) and their average (in *black*). **B:** Same as *A* for blocks where monkeys expected to lose 0, 1, or 3 tokens for incorrect choices. **C:** Same as B for blocks where monkeys expected to win 1, 2, or 3 tokens for correct choices.

**Supplementary Figure 3.**
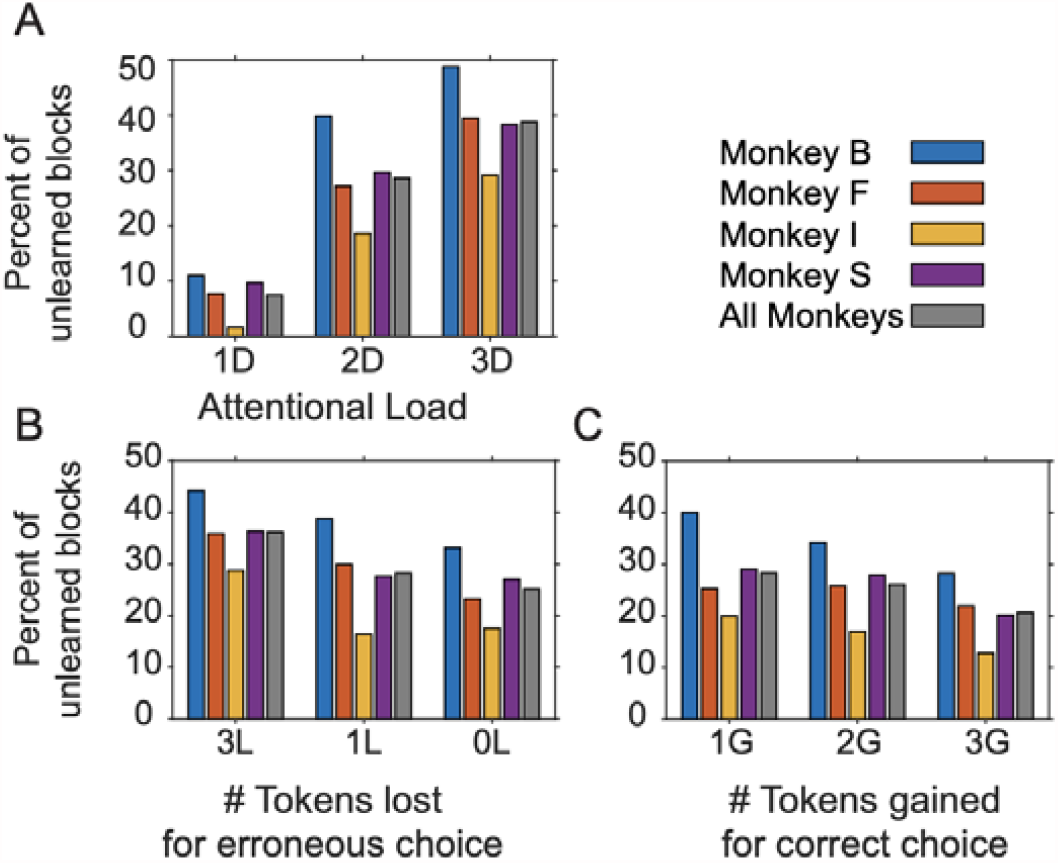
Proportion of unlearned blocks across conditions. **A**: The number of unlearned blocks (*y-axis*) for low/medium/high attentional load (*x-axis*) for each monkey (in *color*) and their average (in *grey*). **B,C:** Same as *A* for loss conditions (*B*) and for the gains conditions (*C*).

